# Self-association configures the NAD^+^-binding site of plant NLR TIR domains

**DOI:** 10.1101/2021.10.02.462850

**Authors:** Hayden Burdett, Xiahao Hu, Maxwell X. Rank, Natsumi Maruta, Bostjan Kobe

## Abstract

TIR domains are signalling domains present in plant nucleotide-binding leucine-rich repeat receptors (NLRs), with key roles in plant innate immunity. They are required for the induction of a hypersensitive response (HR) in effector-triggered immunity, but the mechanism by which this occurs is not yet fully understood. It has been recently shown that the TIR domains from several plant NLRs possess NADase activity. The oligomeric structure of TIR-containing NLRs ROQ1 and RPP1 reveals how the TIR domains arrange into an active conformation, but low resolution around the NAD^+^ binding sites leaves questions unanswered about the molecular mechanisms linking self-association and NADase activity. In this study, a number of crystal structures of the TIR domain from the grapevine NLR RUN1 reveal how self-association and enzymatic activity may be linked. Structural features previously proposed to play roles involve the “AE interface” (mediated by helices A and E), the “BB-loop” (connecting β-strand B and helix B in the structure), and the “BE interface” (mediated by the BB-loop from one TIR and the “DE surface” of another). We demonstrate that self-association through the AE interface induces conformational changes in the NAD^+^-binding site, shifting the BB-loop away from the catalytic site and allowing NAD^+^ to access the active site. We propose that an intact “DE surface” is necessary for production of the signalling product (variant cyclic ADPR), as it constitutes part of the active site. Addition of NAD^+^ or NADP^+^ is not sufficient to induce self-association, suggesting that NAD^+^ binding occurs after TIR self-association. Our study identifies a mechanistic link between TIR self-association and NADase activity.

## Introduction

Plants possess a suite of intracellular receptors that are able to detect and respond to virulence factors secreted by pathogens into the host cell. Nucleotide-binding leucine-rich repeat receptors (NLRs) directly or indirectly detect effector proteins from a range of different pathogens and activate an effector-triggered immunity (ETI) response. ETI is often characterised by localised cell death around the site of infection, known as the hypersensitive response (HR).

The activation of plant NLRs involves the oligomerization into a resistosome complex (Wang et al., 2019a, Wang et al., 2019b, Burdett et al., 2019, Martin et al., 2020, Ma et al., 2020). Typical NLRs are comprised of three distinct domains. The C-terminal leucine-rich repeat (LRR) domain is involved in effector sensing, autoinhibition of NLR signalling, and facilitating the stabilisation of the oligomeric resistosome (Wang et al., 2019a, Wang et al., 2019b, Martin et al., 2020, Ma et al., 2020). The NB-ARC (nucleotide binding-shared between APAF-1, some plant R proteins and CED-4) domain binds nucleotides (Tameling et al., 2002, Tameling et al., 2006, Williams et al., 2011, Bernoux et al., 2016, Martin et al., 2020, Ma et al., 2020) and is responsible for conformational changes that drive the oligomerisation of the resistosome (Wang et al., 2019a, Wang et al., 2019b, Burdett et al., 2019). The N-terminal domain of NLRs is variable and is used to distinguish different classes of NLRs. Found predominantly in monocots, CC-NLRs possess an N-terminal coiled-coil (CC) domain, postulated to initiate HR upon oligomerisation through plasma membrane association and calcium transport (Wang et al., 2019a, Bi et al., 2021). TIR-NLRs, absent in monocot species, possess an N-terminal Toll/interleukin-1 receptor/resistance protein (TIR) domain, recently shown to function through the enzymatic cleavage of NAD^+^ (Wan et al., 2019, Horsefield et al., 2019). This enzymatic activity is implicated in the production of the putative signalling molecule v-cADPR, which is a biomarker for pathogen resistance (Wan et al., 2019), and potentially involved in ETI (Wan et al., 2019, Duxbury et al., 2020); however, the structure and precise role of this molecule is not clear.

TIR domains also need to self-associate in order to intiate HR signalling (Bernoux et al., 2011, Williams et al., 2014, Schreiber et al., 2016, Nishimura et al., 2017, Zhang et al., 2017, Martin et al., 2020, Ma et al., 2020). Accordingly, there is evidence to suggest that self-association is required for NAD^+^ cleavage (Wan et al., 2019, Horsefield et al., 2019). The structures of the ROQ1 and RPP1 TIR-NLR resistosomes confirm that oligomerization is required for NLR activation, and for the first time show how TIR domains are arranged in an activated NLR (Martin et al., 2020, Ma et al., 2020). The TIR domains of both RPP1 and ROQ1 form a tetrameric assembly atop the NB-ARC domains, in a “dimer of dimers” arrangement. This conformation is facilitated by two key interfaces, the “AE interface” and the “BE interface” (Figure 1A,B).

**Figure 1.**
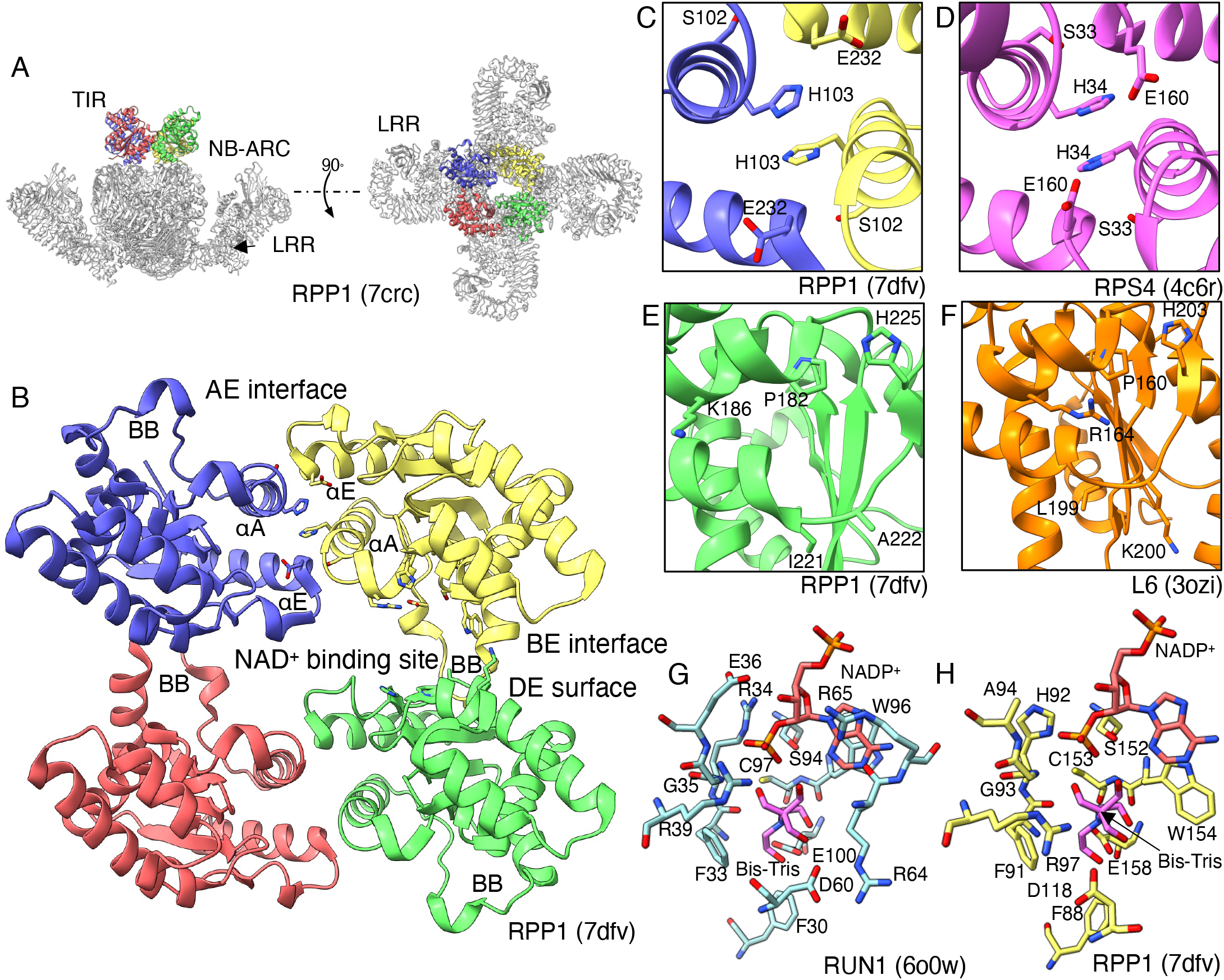
The TIR-domain tetramer, with key interfaces and binding sites. **A**, relative position and conformation of the RPP1^TIR^ tetramer in relation to the RPP1 resistosome (PDB ID: 7crc). **B**, locations of interfaces and binding sites are shown on the RPP1^TIR^ tetramer (PDB ID: 7dfv). **C** and **D**, residues involved in formation of the AE interface, of RPP1^TIR^ and RPS4^TIR^ (PDB ID: 4c6r), respectively. **E** and **F**, residues on the DE surface of RPP1^TIR^ and L6^TIR^ (PDB ID: 3ozi), respectively. **G** and **H**, residues in the Bis-Tris/NADP^+^ binding site of RUN1^TIR^ (PDB ID: 6o0w) and RPP1^TIR^, respectively.

The AE interface (mediated by α-helices A and E), a consistent feature found in crystal structures of plant NLR TIR domains (Williams et al., 2014, Williams et al., 2016, Zhang et al., 2017, Nishimura et al., 2017), is also found in the arrangement of the TIR domains in the ROQ1 and RPP1 resistosomes (Martin et al., 2020, Ma et al., 2020). The hallmark of the AE interface is a pair of intercalaticing histidines at the core of the interface, surrounded by supporting interactions from adjacent aspartate, glutamate and serine residues (Williams et al., 2014) (Figure 1B-D). A DE interface had also been proposed as a possible secondary interface in plant TIR domains, although it has been observed less frequently in crystal structures, shows substantial variability between different structures, and mutations of corresponding residues have less effects on self-association and cell death, compared to the AE interface (Zhang et al., 2017, Nishimura et al., 2017). The ROQ1 and RPP1 resistosome structures demonstrate that the corresponding DE surface is important for NLR function, but as part of an asymmetric interface with the BB loop and the NAD^+^ binding site of another TIR monomer, forming the BE interface (Figure 1B,E-F). An alanologous arrangement of TIR domains, facilitated by the AE and BE interfaces, is observed in the crystal structure of the TIR-domain NADase from the mammalian protein SARM1, which is involved in axon degeneration (Horsefield et al., 2019).

While there is no question of the significant advancement made by the ROQ1 and RPP1 structures, the lower resolution of the TIR domains, absence of NAD^+^ or products bound, and no inactive resistosome to compare and contrast, leave some questions unanswered about the activation and NAD^+^ binding of the TIR domains in the resistosome. While the mechanism of NAD^+^ hydrolysis or v-cADPR formation is not clear, key residues involved in NADase activity have been identified in the crystal structure of grapevine TIR-NLR RUN1 (Horsefield et al., 2019), many of which are conserved in ROQ1 and RPP1 (Figure 1G,H), and in many other plant TIRs (Figure S1). At the base of this NAD^+^ binding pocket is a catalytic glutamate residue, and surrounding the surface of the pocket are numerous aromatic and positively charged residues, thought to coordinate the ribose, nicotinamide, adenosine and phosphate groups (Figure 1G,H). By analysing multiple structures of RUN1^TIR^ and comparing them to the ROQ1 and RPP1 resistosomes, we propose a structural mechanism by which self-association activates NAD^+^-cleavage activity. We show that the formation of the AE interface results in conformational changes in the NAD^+^-binding site, resulting in the movement of the BB-loop and exposure of the catalytic glutamate residue. RUN1^TIR^ structures without an AE interface have a closed NAD^+^-binding site, preventing enzymatic activity. Our analysis also suggests an intact DE surface is required to facilitate oligomerisation of AE and BE interface-mediated TIR tetramers, and to complete the NAD^+^-binding site.

## Experimental procedures

### Protein production and purification

The cDNA of RUN1^TIR^ (residues 23-198) was inserted into the expression vector pMCSG7 by ligation-independent cloning. Mutations were introduced into this construct using the Q5® Site-Directed Mutagenesis Kit (New England Biolabs), and proteins were expressed in *Escherichia coli* BL21 (DE3) cells, using the autoinduction method (Studier, 2005). Overnight cultures were used to seed 1 L autoinduction media cultures containing 100 μg.mL^-1^ ampicillin in 2.5 L baffled flasks. Cultures were grown at 37°C to OD_600_ of 0.6-0.8, before the temperature was lowered to 15°C. Cells were grown for an additional 12-18 h before being harvested by centrifugation. Cells containing RUN1^TIR^ were resuspended in a lysis buffer containing 50 mM HEPES pH 8.0, 500 mM NaCl, 30 mM imidazole, 1 mM DTT and 1 mM PMSF. Cells were lysed by sonication, and the lysate was clarified by centrifugation at 39,191 x *g* for 45 min at 4°C.

Clarified lysate was applied to a 5 mL HisTrap FF column (Cytiva) pre-equilibrated with 50 mM HEPES pH 8.0, 500 mM NaCl and 30 mM imidazole. The column was washed with the lysis buffer to remove *E. coli* proteins. RUN1^TIR^ were eluted using a block elution of lysis buffer containing 250 mM imidazole and fractions containing RUN1^TIR^ were pooled. To remove the 6xHis-tag, imidazole was dialysed out using 10,000 MWCO SnakeSkin™ dialysis tubing (Thermo Scientific), against a buffer containing 10 mM HEPES pH 7.5, 150 mM NaCl, 0.5 mM EDTA and 1 mM DTT. The 6xHis-tag of RUN1^TIR^ was removed by incubation with 6xHis-tagged tobacco etch virus (TEV) protease overnight at 4°C.

Cleaved RUN1^TIR^ was reapplied to the HisTrap column to remove the TEV protease and contaminants. The sample was then applied to a Superdex S75 26/600 column (Cytiva) pre-equilibrated with 10 mM HEPES pH 7.5, 150 mM NaCl and 1 mM DTT. Peak fractions were pooled and concentrated to 10 mg.mL^-1^ using a 10,000 MWCO centrifugal concentrator (Merck Millipore). Protein was flash-frozen in liquid nitrogen and kept at −80°C until required. All RUN1^TIR^ mutants were purified in the same manner as the wild-type protein.

### Crystallisation, X-ray data collection and crystal structure determination

Initial crystallization screening was performed at 293 K, using 200 nL drops in 96-well drops, at a concentration of 10 mg.mL^-1^. For co-crystallisation with ligands, the protein was incubated with a range of ligands, at a range of concentrations. Small crystals appeared after 3 days in multiple conditions.

“RUN1^TIR^ ΔAE” crystals were produced by the hanging-drop vapour diffusion method, with 0.8 μL of protein (10 mg.mL^-1^) and 0.8 μL of well solution (0.1 M sodium acetate pH 4.6, 1.8 M ammonium sulfate), and appeared within 4-5 days. Crystals were flash-cooled in liquid nitrogen, using reservoir solution with 10% w/v glycerol and 10% w/v ethylene glycol as a cryo-protectant.

“RUN1^TIR^ RPV1-like” and “RUN1^TIR-E100A^ RPV1-like” crystals were produced using the hanging-drop method, with drops containing 1 μL of protein (10 mg.mL^-1^), and 1 μL of well solution (0.2 M Lithium Sulfate, 0.1 M HEPES pH 7.0, 20% PEG3350), and appeared within 7-10 days. Crystals were added to cryoprotectant (0.2 M Lithium Sulfate, 0.1 M HEPES pH 7.0, 20% PEG3350, 10% w/v glycerol, 10% w/v ethylene glycol and 10 mM NAD^+^) and soaked for 2-5 min, before being flash-cooled in liquid nitrogen.

“RUN1^TIR-E100A^ NAD^+^” crystals were produced using the hanging-drop method, with drops containing 1 μL of protein (10 mg.mL^-1^), 1 μL well solution (2 M ammonium sulfate 0.1 M TrisBase pH 8.75), and 1 μL 30 mM NAD^+^, and appeared after ~7 days. Crystals were added to cryoprotectant (2 M ammonium sulfate, 0.1 M Tris pH 8.75, 20% w/v glycerol, 30 mM NAD^+^) and soaked for 30 seconds, before being flash-cooled in liquid nitrogen.

### Structure analysis

Data collection was performed using the Blu-Ice software, indexed and integrated using XDS (Kabsch, 2010) and/or DIALS (Beilsten-Edmands et al., 2020) and scaled using AIMLESS (Evans, 2006). The “RUN1^TIR^ ΔAE”, “RUN1^TIR^ RPV1-like” and “RUN1^TIR-E100A^ NAD^+^” structures were determined by molecular replacement using Phaser (McCoy, 2007), with the structure of “RUN1^TIR^ NADP^+^” (PDB ID 6o0w; (Horsefield et al., 2019)) as a template. Refinement was performed iteratively using phenix.refine (Adams et al., 2013) and Coot (Emsley et al., 2010), and structures were analysed using PISA (Krissinel and Henrick, 2007), Pymol (Schrodinger) and ChimeraX (Goddard et al., 2018). Figures were prepared using ChimeraX.

### SEC-MALS experiments

SEC-MALS (size-exclusion chromatography coupled with multi-angle light scattering) was performed as described by Horsefield et al. (2019), using a Superdex S200 1/150 Increase column (Cytiva). RUN1^TIR^ was incubated at room temperature with 2 mM NAD^+^ or NADP^+^ for 30 minutes prior to loading onto the column.

### Fluorescence assay for NAD^+^ hydrolysis

Assays using a fluorescent NAD^+^ analog, εNAD (nicotinamide 1,N^6^-ethenoadenine dinucleotide), were performed as described by Pergolizzi et al. (2011) and Horsefield et al. (2019). Briefly, purified 50 μM RUN1^TIR^, and mutants were added to a black 96 well plate in 10 mM HEPES pH 7.5, 150 mM NaCl with 25% (w/v) PEG3350. After incubation in the dark for 15 min at 25°C, 100 μM of εNAD was added to each well. Fluorescence intensity was measured in a CLARIOstar® microplate reader, using an excitation of wavelength of 310-330 nm and emission wavelength of 390-410 nm, with readings every two minutes over four hours. The change in fluorescence over time was determined by calculation of slopes of the linear component of the curves. Data analysis was performed using Prism (GraphPad). All measurements, including controls were made in triplicate.

### Cycling assays for NAD^+^ hydrolysis

Cycling assays were performed as described by (Horsefield et al., 2019), except using purified protein instead of bacterial lysate extracts. To prepare the NAD^+^ samples, 100 μM of RUN1^TIR^ was mixed with 10 μL of a 50% slurry of Ni-NTA beads washed in 10 mM HEPES pH 7.5, 150 mM NaCl, and NAD^+^ at various concentrations (100-5000 μM) in a final volume of 200 μL. At set time points (ranging from 20 minutes, to up to 24 hours), 20 μL was removed and heated at 95°C for one minute to deactivate the protein, then stored at −20°C until required. This sample was then used in the enzyme-linked cycling assay, and NAD^+^ concentrations of the samples were determined using a standard curve of known NAD^+^concentrations.

## Results

### RUN1^TIR^ readily crystallises under several different conditions

For many TIR-NLRs, recombinant production of the full-length protein has been challenging, but it has been possible in *E. coli* for the TIR domain only. Bacterially expressed RUN1^TIR^ yielded crystals in several conditions and we determined four unique structures to date. The “RUN1^TIR^ NADP^+^” crystal structure (6o0w) was the first to be determined and has been described previously (Horsefield et al., 2019). Each of the other RUN1^TIR^ structures have different packing arrangements (Supplementary Figure 2), and when compared to other TIR structures, provide insights into how plant TIR domains might self-associate during the processes of regulation and activation. Interestingly, none of the structures features both the AE and BE interfaces observed in the ROQ1 and RPP1 resistosome structures. Three structures of note will be discussed further here: “RUN1^TIR^ ΔAE”, lacking the AE interface (as well as the symmetric DE interface) in the crystal packing; “RUN1^TIR^ RPV1-like” that features the AE interface and has a packing arrangement similar to the RPV1^TIR^ crystal structure (PDB ID:5ku7) (Williams et al., 2016); and “RUN1^TIR-E100A^ NAD^+^”, a structure of RUN1 with a mutation to the catalytic glutamate, bound to NAD^+^. Each structure informs how AE interface formation, and of NAD^+^ binding to the BE interface may occur in context of the full-length resistosome structures of ROQ1 and RPP1 (Figure 2A).

**Figure 2.**
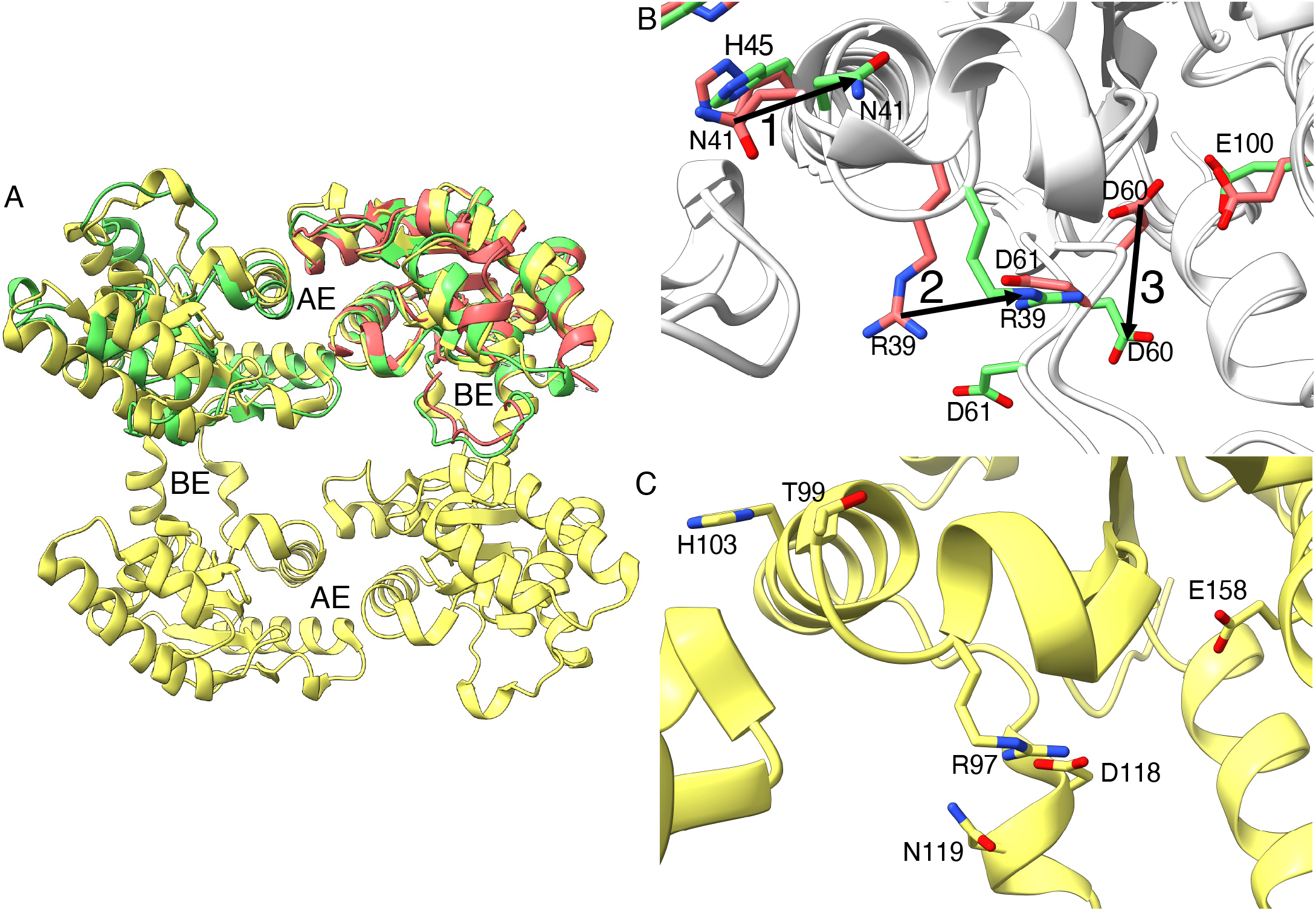
AE interface formation opens BB loop. **A**, superimposition of the RPP1^TIR^ tetramer from the resistosome (yellow, PDB ID: 7dfv), the RUN1^TIR^ structure with no AE interface (red, PDB ID: 7rts) and RUN1 structure with the AE interface (green, PDB ID: 7rx1). **B**, key residues involved in confirmational changes between different RUN1^TIR^ structures. The steps involve: (1) residues at the N-terminal end of the αA helix need to move upon formation of AE interface in RUN1^TIR^. This movement causes (2) a conserved arginine residue to move toward (3) a conserved aspartate at the C-terminal end of the βB sheet. The arginine moves the conserved aspartate away from the conserved catalytic glutamate, partially opening the proposed NAD^+^ binding site. The formation of the BE interface then follows, creating the TIR-domain tetramer, and the complete NAD^+^ binding site. **C** The open conformation can be seen in the RPP1^TIR^ tetrameric structure.

### AE interface formation induces conformational changes in the RUN1^TIR^ NAD^+^-binding site

When RUN1^TIR^ structures that possess an AE interface are compared with those that do not, distinct structural changes in positions of NAD^+^-binding-site residues become evident. All RUN1^TIR^ structures contain the canonical AE interface, except for the “RUN1^TIR^ ΔAE” crystal structure. The histidine (H45) in the core of the AE interface is rotated approximately 90° between the two structures. On the N-terminal end of the αA helix, there is a cis-peptide conformation between residues 40 and 41 that is present in the “RUN1^TIR^ ΔAE” structure, but not in other crystals. This conformation places the “RUN1^TIR^ ΔAE” N41 side chain away from the β-sheet core of the TIR domain (Figure 2B).

If N41 were to adopt this cis-peptide conformation in a RUN1 crystal with an AE interface, there would be a steric clash between it and E175 from the other TIR in the dimer. Superposition of a “RUN1^TIR^ ΔAE” crystal monomer onto one of the molecules of the AE-interface dimer pair from the “RUN1^TIR^ RPV1-like” crystal reveals a number of additional clashes at the interface, including R39-Y174, H45-S176, and F40-G173 (considering the “RUN1^TIR^ ΔAE” and the “RUN1^TIR^ NADP^+^” monomers in the superposition, respectively). The rest of the AE interface is clash-free; thus, if R39, F40, N41 and H45 shift to accommodate G173, Y174, E175 and S176, the AE interface can form. If the RUN1^TIR^ domains were brought together in the confirmation seen in ROQ1 and RPP1 resistosome structures, the N41 shifting away would lead to a cis-trans isomerisation of the peptide. This peptide isomerisation would in turn move R39 and F40 away from the AE interface, and toward the NAD^+^-binding site. Non-proline cis-trans peptide isomerisation is often found to play roles in the regulation of protein activities, including many signalling proteins (Joseph et al., 2012).

Movement of this conserved argininine (R39) in RUN1, in response to AE-interface formation, is likely key to conformational changes in the binding site. Superposition of the “RUN1^TIR^ ΔAE” structure “RUN1^TIR^ RPV1-like” crystals AE-interface dimers shows a large clash between R39 of RUN1^TIR^ from the AE-interface dimer and D61 from the “RUN1^TIR^ ΔAE” structure. D60 and D61 are moved out to accommodate R39 in the AE-interface dimer (Figure 2B). Another substantial change, when comparing the “RUN1^TIR^ RPV1-like” and “RUN1^TIR^ ΔAE” crystals, involves D60 and D61. In the “RUN1^TIR^ ΔAE” crystals, D60 is in close proximity to E100 (Figure 2B). D61 is also positioned closer to E100 in the NAD^+^-binding site, compared to the “RUN1^TIR^ RPV1-like crystals. D60 and D61 move away from E100 upon AE-interface formation (Figure 2B). This change in position of D60 effectively ‘opens’ the NAD^+^-binding site, as it enables the whole BB loop to move further away from the core of the TIR domain, increasing solvent accessibility. This open state can be observed in the RPP1 resistosome structure (Figure 2C). The opening can be measured by the distance of D60 to E100 side chain atoms. In the RUN1 closed conformation, D60 carboxyl groups are within 3 □ of E100, while in the open confirmation, they move to 6-7 □ away. These distances are even greater for the RPP1 resistosome structure. We propose that a combination of the F40-N41 peptide isomerisation, and the steric clashes that would occur between E175 and H45, and Y174 and R39, drive the opening of the NAD^+^-binding site, by shifting D60 and D61 away from the catalytic glutamate (Figure 2B). This initial opening of the binding site would be required to allow the BB-loop to be positioned under the DE surface in the BE dimer, as seen in both ROQ1 and RPP1 structures.

### DE surface represents one half of the BE interface and forms part of the NAD^+^-binding site

The AE interfaces are highly similar in different plant TIR-domain structures, but this is not the case for the DE interfaces. In all AE-interface containing plant TIR structures, the two subunits have identical orientations relative to one another. By contrast, the orientations of the subunits in the DE-interface dimers vary significantly, even between different RUN1^TIR^ structures. Mutations in the DE interface appear have less impact on HR and self-association than those in the AE interfaces (Bernoux et al., 2011, Bernoux et al., 2016, Zhang et al., 2017). The ROQ1 and RPP1 resistosome structures reveal that the DE interface surface in fact forms one half of the BE interface, and not a symmetric interface as previously expected.

During attempts to crystallise RUN^TIR^ with NAD^+^ bound, we noticed multiple RUN1^TIR^ structures had NAD^+^ or the non-hydrolysable analogue carbaNAD binding at the DE surface (Figure 3A,B). The ROQ1 and RPP1 resistosome structures reveal that this NAD^+^ molecule would sit in the open cleft of the BE interface at the active site of the neighboring monomer. Strikingly, it occupies the same position as the ATP molecule captured in the RPP1 and ROQ1 TIR domains (Figure 3A-C). There was some additional density in this structure around the catalytic binding site, particulary around reisude W96, that we could not definitively build a model into; however, there was clear density for NAD^+^ bound to the DE surface of the BE interface (Figure 3D).

**Figure 3.**
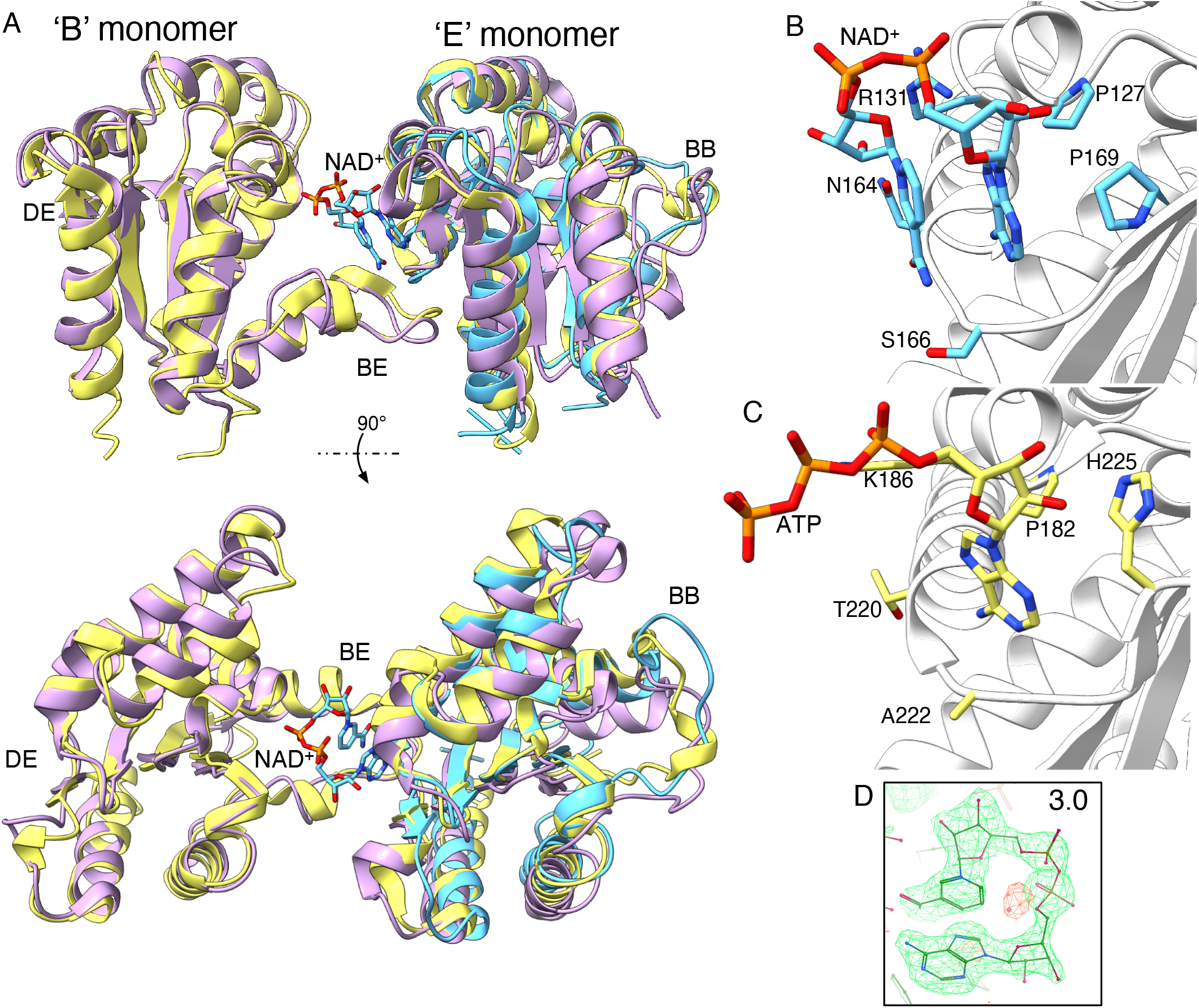
NAD^+^ bound to RUN1^TIR^ may represent biologically the relevant NAD^+^-binding site of plant TIRs. **A**, a superposition of RUN^TIR-E100A^ (cyan, PDB ID: 7s2z) onto the ‘E’ monomer of the ROQ1 (purple, PDB ID: 7jlx) and RPP1 (yellow, PDB ID: 7dfv), BE interface dimer. NAD^+^ sits directly above the BB loop, with the nicotinamide moiety positioned toward the open primary NAD^+^ binding site in the ‘B’ monomer. **B** and **C** shows residues of the DE surface of RUN1^TIR^ and RPP1^TIR^ in proximity to the bound NAD^+^ and ATP, respectively. **D** shows the omit map of the NAD^+^ molecule from RUN^TIR-E100A^ structure in **A** and **B**.

The NAD^+^ molecule folds back onto itself, with the nicotinamide group packing against the adenine moiety (Figure 3D). The adenosine moiety of NAD^+^ makes contact with the DE surface, leaving the nicotinamide moiety free to move toward the catalytic glutamate of the NAD^+^ active site. Residues P127, R131, A163, N164, L165, S166 and P169 are all involved in interactions or contribute to a large proportion of the buried surface area of this binding site (Figure 3B). Mutations of three of these residues in other plant TIRs impair HR; the mutation of the residue equivalent to P127 impairs HR in L6 (Bernoux et al., 2011, Bernoux et al., 2016), N (Dinesh-Kumar et al., 2000, Mestre and Baulcombe, 2006), RPS4 (Swiderski et al., 2009) and RPV1 (Williams et al., 2016); the mutation of the residue equivalent to R131 impairs HR in L6 (Bernoux et al., 2011, Bernoux et al., 2016), RPS4 (Swiderski et al., 2009), RPV1 (Williams et al., 2016) and SNC1 (Zhang et al., 2017); and the mutation of the residue equivalent to S166 impairs HR in L6 (Bernoux et al., 2011, Bernoux et al., 2016) and RPS4 (Swiderski et al., 2009). While they all impair HR, they do not all affect self-association of L6 (Bernoux et al., 2011, Bernoux et al., 2016, Zhang et al., 2017).

### NAD^+^ and NADP^+^ do not induce self-association of RUN1^TIR^ in solution

Based on the differences in the active-site accessibility between the RUN1^TIR^ structures with and without the AE interface, we wanted to test, using the SEC-MALS technique, if NAD^+^binding could stimulate AE interface-mediated self-association. Purified RUN1^TIR^ protein was incubated with NAD^+^ or NADP^+^, prior to running SEC-MALS. Given that RUN1^TIR^ is monomeric in solution, an increase in molecular weight after incubation with NAD^+^ would imply that NAD^+^binding stimulates RUN1^TIR^ self-association. At a range of protein concentrations, 2 mM of NAD^+^ or NADP^+^ do not stimulate RUN1^TIR^ to self-associate (Figure 4 and Supplementary Figure 3). RUN1^TIR(RRAA)^ is a mutant with higher NADase activity, presumably due to a more accessible NAD^+^ binding site (Horsefield et al., 2019). This mutant is monomeric in solution at 2 mg.mL^-1^, and addition of 2 mM NAD^+^ or 2 mM NADP^+^also does not stimulate self-association or higher-order oligomers. Therefore, we suggest that in the context of full-length plant NLRs, self-association of the NB-ARC domain and resistosome formation facilitates AE-interface and BE-interface formation. These results, along with the closed BB-loop conformation of RUN1 TIR domains without an AE interface suggests the AE and BE oligomerisation precedes NAD^+^ or NADP^+^ binding. Further experiments would be required to test this hypothesis, however.

**Figure 4.**
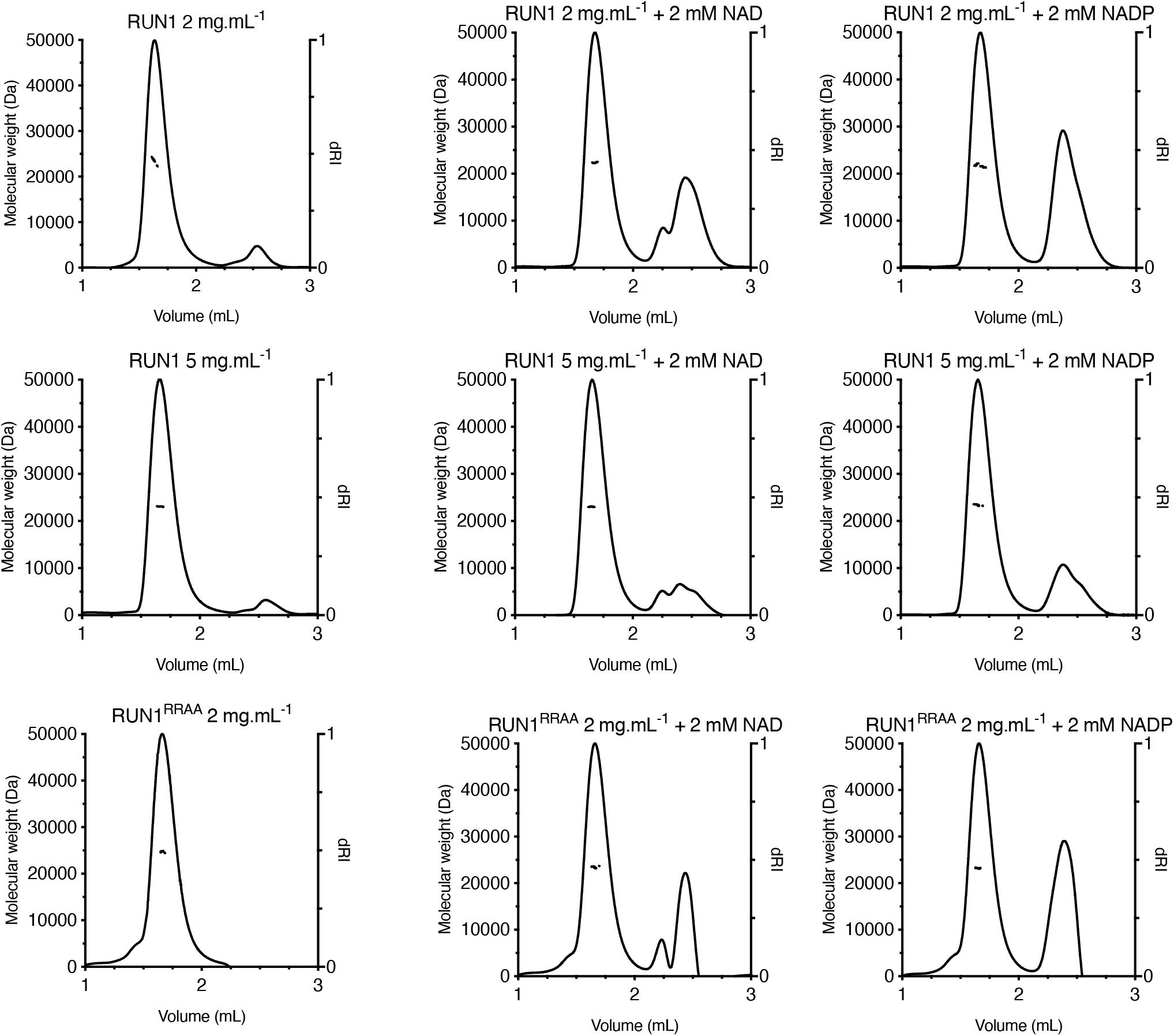
NAD^+^ and NADP^+^ do not induce self-association of RUN1^TIR^ or RUN1^TIR(RRAA)^ proteins. The proteins were incubated with 2 mM NAD^+^ or 2 mM NADP^+^ for 20 minutes at room temperature, before SEC-MALS analysis. RUN1^TIR^ at 2 and 5 mg.mL^-1^ elutes with a peak of a molar mass consistent with a monomeric RUN1^TIR^ (~ 23 kDa). Addition of 2 mM NADP^+^ or NADP^+^ does not stimulate an increase in this molecular weight at 2 or 5 mg.mL^-1^. The mutant RUN1^TIR(RRAA)^ with increased activity was also tested at 2 mg.mL^-1^ protein concentration, and similar results were observed. NAD^+^ and NADP^+^ can also be seen on the differential refractive index (dRI) trace, eluting at ~ 2.5 mL. RUN1 in this instance refers to the TIR domain of RUN1, NAD to NAD^+^ and NADP to NADP^+^.

### Effects of mutations, in the DE-surface NAD^+^-binding site, on NADase activity

The effects of mutations in the AE interface and the DE surface, as well as the NAD^+^-binding site, on the NAD^+^-cleavage activity, were tested by the fluorescence NAD^+^ assay (Horsefield et al., 2019). As seen in Figure 5A, R131E, the mutation of a residue on the periphery of the BE interface does not impair NADase activity, while P169R, closer to the proposed active site, reduces but does not abolish NADase activity. F33A, R34A, S94A, W96A, W96T and E100A mutations, all within or near the NAD^+^-binding site, impair NAD^+^-cleavage activity, consistent with previous results. R39A, a mutation of a residue important for the AE interface-induced switch, knocks out NADase activity completely. D44A, R64A and C97A all severely reduce NADase activity. Consistent with the need to self-associate, we observe an increase in the ability of purified RUN1^TIR^ to cleave NAD^+^, when immobilised on Ni-NTA beads (Figure 5B).

**Figure 5.**
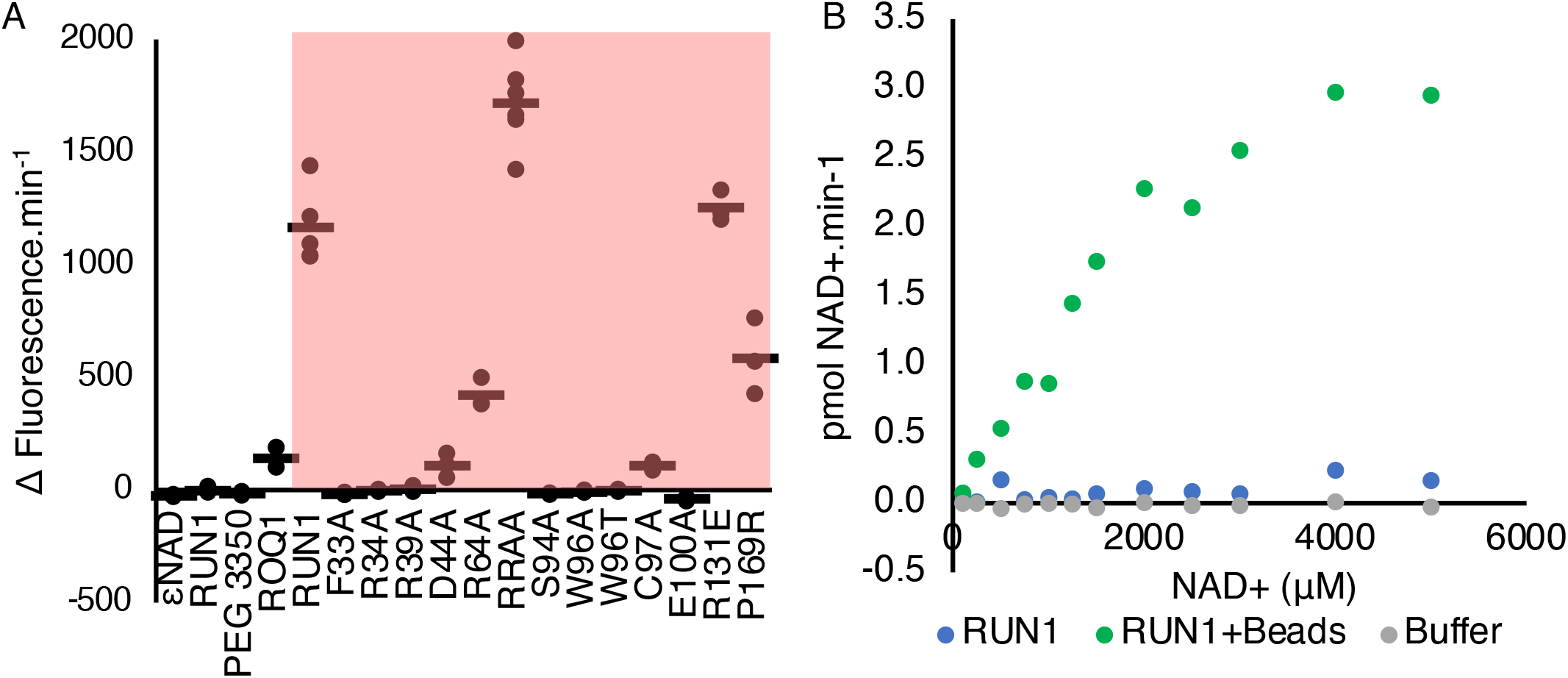
NADase activity of RUN1 TIR domains requires self-association, and intact NADase binding site. **A**, Fluorescence-based NADase assay with the molecular-crowding agent PEG3350 added. Samples in red shaded area contain PEG 3350 as a molecular crowding agent **B**, NAD^+^ assay, using NAD^+^ as a substrate, His-tagged protein on Ni-NTA agarose beads, and the cycling assay to measure NAD^+^ concentration.

## Discussion

### AE interface is required for NADase activity

We show that conserved residues of plant TIR domains are involved in a series of conformational changes, regulating NAD^+^ binding site accessability and BE interface formation. Mutations that prevent formation of the AE interface impair self-association (Williams et al., 2014, Zhang et al., 2017), HR (Williams et al., 2014, Williams et al., 2016, Zhang et al., 2017, Martin et al., 2020, Ma et al., 2020), NADase activity (Wan et al., 2019) and variant cyclic ADPR (v-cADPR) formation (Wan et al., 2019). NAD^+^ does not induce self-association of TIR domains (Figure 4); thus, we propose that the AE interface must form to allow NAD^+^ to bind.

Based on structural information, a key residue that transfers the AE interface formation into BB-loop opening and the formation of the active site in RUN1, is R39 (Figure 2B). In different plant TIR domains, mutation of this residue does not substantially affect TIR-domain self-association (Bernoux et al., 2011); however, the mutation impairs NADase activity (Figure 5), autoactivity by TIR domains (Bernoux et al., 2011, Williams et al., 2016) and effector-dependent HR (Mestre and Baulcombe, 2006). This residue moves toward the BB-loop upon AE interface formation, facilitating the opening of the BB-loop. This is likely occurring around residues D60 and D61 in RUN1 (Figure 2B). D60 in particular is highly conserved among plant TIRs (Figure S1). A cis-to-trans peptide isomerisation of F40-N41 likely facilitates the movement of R39; however, experimental evidence in solution would be required to confirm this. Thus, formation of the AE interface acts as an allosteric regulator of NADase activity, with R39 propagating this signal from AE interface to NAD^+^ active site. This additional layer of regulation of NADse activity likely helps prevent cell death or immune sigalling in the absence of pathogens.

### DE surface may facilitate v-cADPR formation

The ROQ1 and RPP1 resistosome structures provide new insight into the DE surface mutations tested in the past. In contrast to the symmetrical DE interfaces observed in crystal packing of several plant TIR domains, the DE surface forms an asymmetric interface with the BB-loop of an adjoining AE dimer pair in the ROQ1 and RPP1 resistosome structures. Some mutations to the conserved glycine in this interface support this, with mutation to large bulky charged residues having a much more significant effect on HR, self-association and NADase activity than more subtle mutations. For example, RPP1-WsB G223A shows only reduced NADase activity (Ma et al., 2020), while RBA1 G151R, RPS4 G151R, and RPP1-NdA G229R have NADase activity abolished (Wan et al., 2019). Similarly, the L6 G201C and G201Y mutations have only reduced TIR autoactivity (Bernoux et al., 2011), compared to complete loss of autoactivity seen in L6 G201R (Bernoux et al., 2011), RPV1 G161R (Williams et al., 2016), RBA1 G151R, RPS4 G151R, and RPP1-NdA G229R (Wan et al., 2019). The L6 mutants also show a consistent pattern with regards to self-association, with L6 G201C still able to self-associate, L6 G201Y having slightly reduced self-association compared to the wild-type protein, and G201R not being able to self-associate at all (Bernoux et al., 2011). This arginine mutation may potentially protrude into the NAD^+^-binding site, additionally inhibiting NADase activity. This surface of the TIR domain is crucial for both NADase activity and self-assocaiiton of TIR domains, and appears to form one half of the catalytic site.

The RPP1 G223A mutant has only reduced NADase activity, and induces HR (Ma et al., 2020), but the analogous mutation G153A in ROQ1 completely abolishes HR (Martin et al., 2020). Mutation of RPP1 A222E severly abrogates NADase activity, and also HR (Ma et al., 2020). Assuming v-cADPR is the signalling product, disruption of the second half of the active site with a subtle alanine mutation may still allow NAD^+^ cleavage, but disrupt the BE interface such that v-cADPR cannot be produced. We observe some residual NADase activity for proteins with mutations in this surface (R131E and P169R) in RUN1 (Figure 5A). Analysis of the products of such mutations *in vivo* and *in vitro* would provide greater insight to the role of the BE interface in v-cADPR cyclisation.

### RUN1 RRAA mutant

The RUN1^TIR^ RRAA mutant has increased NADase activity compared to the wild-type protein; however, it does not show stronger self-association *in vitro*, as measured by SEC-MALS (Figure 4). This mutant also has increased autoactivity by the TIR domain *in planta*, so much so that it is able to induce HR in *eds1* knock-out lines, much like SARM1 is able to (Horsefield et al., 2019). We suggest the RRAA mutant is able to induce cell death through loss of NAD^+^, as SARM1^TIR^ can when transientially expressed *in planta* (Horsefield et al., 2019). Residues on the BB-loop thus are key to both regulating NADase activity, and to facilitating correct active site formation.

### SARM1 NADase activity

The observations made here on how self-association relates to NAD^+^ cleavage in plant TIR domains are consistent with the observations made for the TIR domain from the mammalian protein SARM1. The crystal structures of SARM1^TIR^ and its mutants have revealed that SARM1^TIR^ also requires two major interfaces, a BB-loop mediated interface, and an AE interface (Figure S4A-C). Mutational studies show that both these interfaces are crucial for NADase activity (Horsefield et al., 2019). SARM1 G601P, a mutation in the BB-loop, locks the TIR into a closed conformation (Figure S4D,E). This mutation also impairs ROQ1 HR (Martin et al., 2020), suggesting this BB loop open and closing mechanism is conserved between SARM1 and plant TIR domains. A SARM1 mutation in the AE interface, involving H685, also impairs activity, by disrupting interactions with Y568, R570 and H685 (Figure S4D,F). This histidine mutation affects the interaction between residues around the position of R39, D60 and D61 in RUN1^TIR^, and suggesting both RUN1 and SARM1 form AE interfaces and change BB-loop conformations in an analogous manner.

### Conclusions

We propose that the AE interface is required to open the NAD^+^-binding site and our structural work reveals the molecular basis for this process. The absence of an AE-interface formation precludes NAD^+^ binding. AE interface-mediated self-association therefore works as a regulatory mechanism, activating TIR domains only when they are positioned correctly. In light of the ROQ1 and RPP1 structures revealing the arrangement of TIR domains on top of the resistosome, we propose that the DE surface is required to facilitate NAD^+^ binding to the BB-loop and packing of the AE-interface TIR domain dimers. The DE surface may also be required to aid correct cyclisation of v-cADPR, the possible signalling product of NAD^+^cleavage. Whether all TIR-containing NLRs form a tetramer when activated is still not known; more structural data will be required to test this hypothesis.

**Table 1.** Data collection and refinement statistics.

| Crystals | RUN1<sup>TIR</sup> ΔAE | RUN1<sup>TIR</sup> RPV1-like | RUN1<sup>TIR(E100A)</sup> NAD<sup>+</sup> |
| --- | --- | --- | --- |
| PDB ID | 7rts | 7rx1 | 7s2z |
| X-ray wavelength | 0.9537 Å | 0.9537 Å | 0.9537 Å |
| Resolution range (Å) | 38.62 - 1.74 (1.80 - 1.74) | 33.49 - 1.89 (1.96 - 1.89) | 47.08 - 2.35 (2.43 - 2.35) |
| Space group | P 41 21 2 | C 1 2 1 | P 2 21 21 |
| Unit cell (Å/°) | 50.37 50.37 120.31<br>90 90 90 | 152.44 41.47 77.23<br>90 118.51 90 | 79.74 116.64 122.22 90 90<br>90 |
| Total reflections | 429,425 (42,566) | 115,763 (11,165) | 350,462 (31,322) |
| Unique reflections | 16,630 (1,600) | 34,127 (3,294) | 47,985 (4,645) |
| Multiplicity | 25.8 (26.6) | 3.4 (3.4) | 7.3 (7.3) |
| Completeness (%) | 99.90 (99.94) | 99.48 (97.60) | 99.80 (98.22) |
| Mean I/sigma(I) | 41.4 (6.2) | 12.1 (0.9) | 13.1 (2.5) |
| Wilson B-factor (Å<sup>2</sup>) | 25.65 | 34.67 | 36.54 |
| R-merge | 0.048 (0.492) | 0.062 (1.262) | 0.096 (0.736) |
| R-meas | 0.049 (0.502) | 0.074 (1.499) | 0.112 (0.792) |
| R-pim | 0.001 (0.096) | 0.040 (0.801) | 0.057 (0.290) |
| CC<sub>1/2</sub> | 1.000 (0.980) | 0.997 (0.358) | 0.999 (0.906) |
| CC* | 1.000 (0.995) | 0.999 (0.726) | 0.999 (0.975) |
| Reflections used in refinement | 16,614 (1,599) | 34,117 (3,291) | 47,818 (4,634) |
| Reflections used for R-free | 851 (91) | 1,627 (152) | 2,350 (213) |
| R-work | 0.2002 (0.2402) | 0.2085 (0.3207) | 0.2177 (0.2710) |
| R-free | 0.2275 (0.3724) | 0.2279 (0.3392) | 0.2547 (0.3477) |
| Number of non-hydrogen atoms | 1,430 | 2,975 | 5,988 |
| Macromolecules | 1,357 | 2,825 | 5,602 |
| Ligands | 20 | 35 | 266 |
| Solvent | 53 | 115 | 120 |
| No. of molecules in the ASU | 1 | 2 | 4 |
| Protein residues | 165 | 344 | 684 |
| RMS (bonds, Å) | 0.006 | 0.003 | 0.003 |
| RMS (angles, °) | 0.89 | 0.55 | 0.57 |
| Ramachandran favored (%) | 95.65 | 97.65 | 97.19 |
| Ramachandran allowed (%) | 3.73 | 2.06 | 2.81 |
| Ramachandran outliers (%) | 0.62 | 0.29 | 0.00 |
| Rotamer outliers (%) | 1.39 | 0.00 | 0.34 |
| Clashscore | 2.95 | 3.73 | 2.09 |
| Average B-factor (Å<sup>2</sup>) | 31.50 | 39.85 | 46.60 |
| Macromolecules | 31.04 | 39.67 | 46.08 |
| Ligands | 53.07 | 58.23 | 59.29 |
| Solvent | 35.06 | 38.83 | 42.40 |

The statistics were calculated using AIMLESS (Evans, 2006) and MolProbity (Chen et al., 2010).
Statistics for the highest-resolution shell are shown in parentheses.
$$R_{\text{work}} / R_{\text{free}} = \frac{\sum_{\text{hkl}} |F_{\text{obs}_{\text{hkl}}} - F_{\text{calc}_{\text{hkl}}|}{\sum_{\text{hkl}} F_{\text{obs}_{\text{hkl}}}}$$ ; $R_{\text{free}}$ was calculated using randomly chosen 563 5% fraction of data that was excluded from refinement.

## Acknowledgments

We acknowledge the use of the University of Queensland Remote Operation Crystallization and X-Ray Diffraction (UQ-ROCX) Facility within the Centre for Microscopy and Microanalysis, and the macromolecular crystallography (MX) beamlines at the Australian Synchrotron, Victoria, Australia.

Molecular graphics and analyses were performed with UCSF ChimeraX, developed by the Resource for Biocomputing, Visualization, and Informatics at the University of California, San Francisco, with support from National Institutes of Health R01-GM129325 and the Office of Cyber Infrastructure and Computational Biology, National Institute of Allergy and Infectious Diseases.

We thank Jeffrey Nanson and Zhenyao Luo (University of Queensland), Thomas Ve (Griffith University) and Simon Williams (Australian National University) for advice on structure refinement and interpretation.

We also thank Adam Bentham (John Innes Centre) and Megan Outram (Australian National University) for helpful discussions.

We thank Thomas Ve and Mark von Itzstein (Griffith University) for the generous gift of carbaNAD.

## Data availability

Coordinates for all structures deposited have been to the Protein Data Bank: RUN1^TIR^ ΔAE, PDB ID: 7rts; RUN1^TIR^ RPV1-like, PDB ID: 7rx1; RUN1^TIR(E100A)^ NAD^+^, PDB ID; 7s2z.

## Supporting information

This article contains supporting information.

## Funding and additional information

The work was supported by the Australian Research Council (ARC; DP160102244 and DP190102526 to B.K.). B.K. is an ARC Laureate Fellow (FL180100109).

---

**Figure S1.**
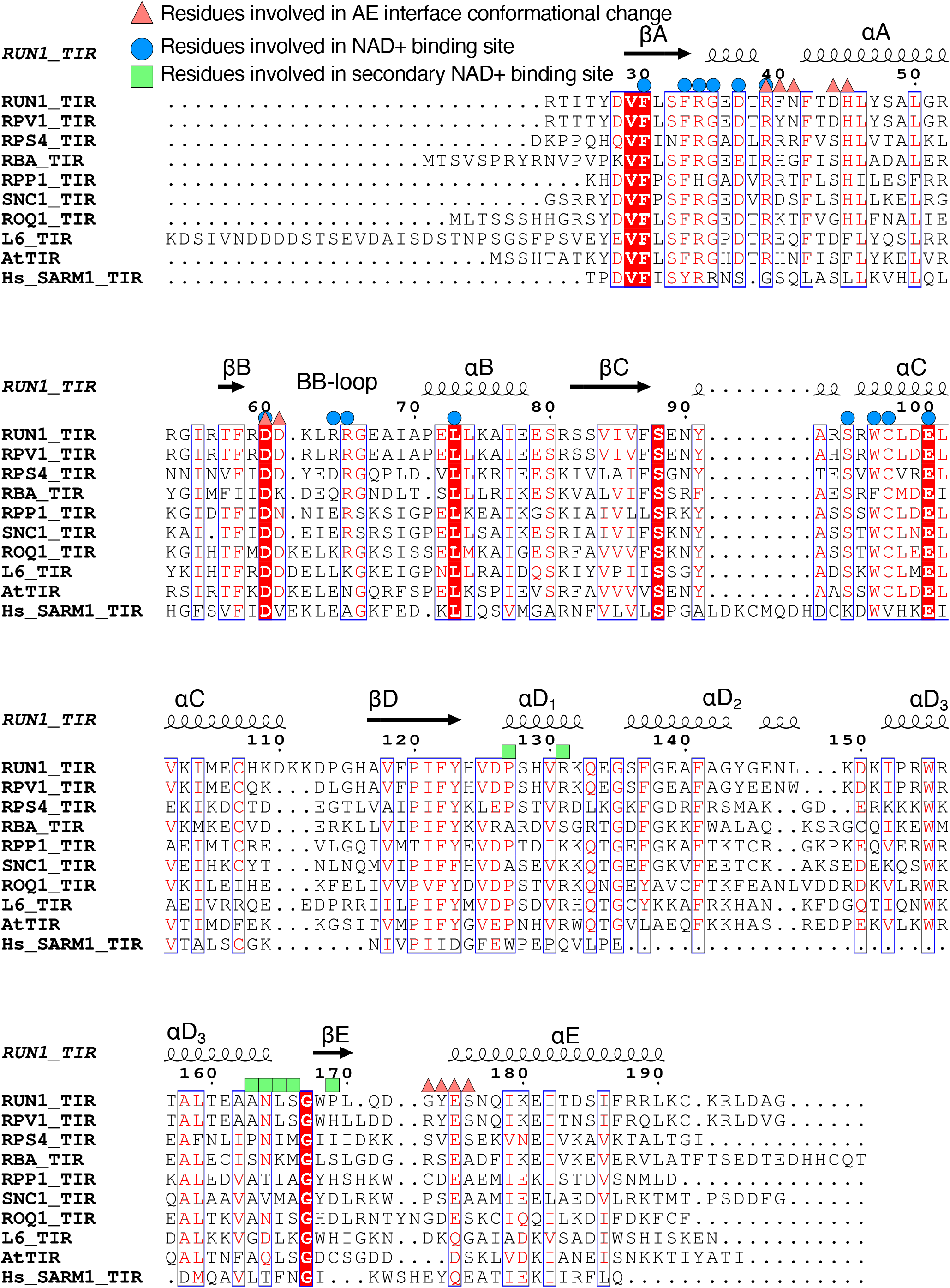
Sequence alignment of various plant TIR domains and SARM1, showing conserved residues important for self-association and NADase activity.

**Figure S2.**
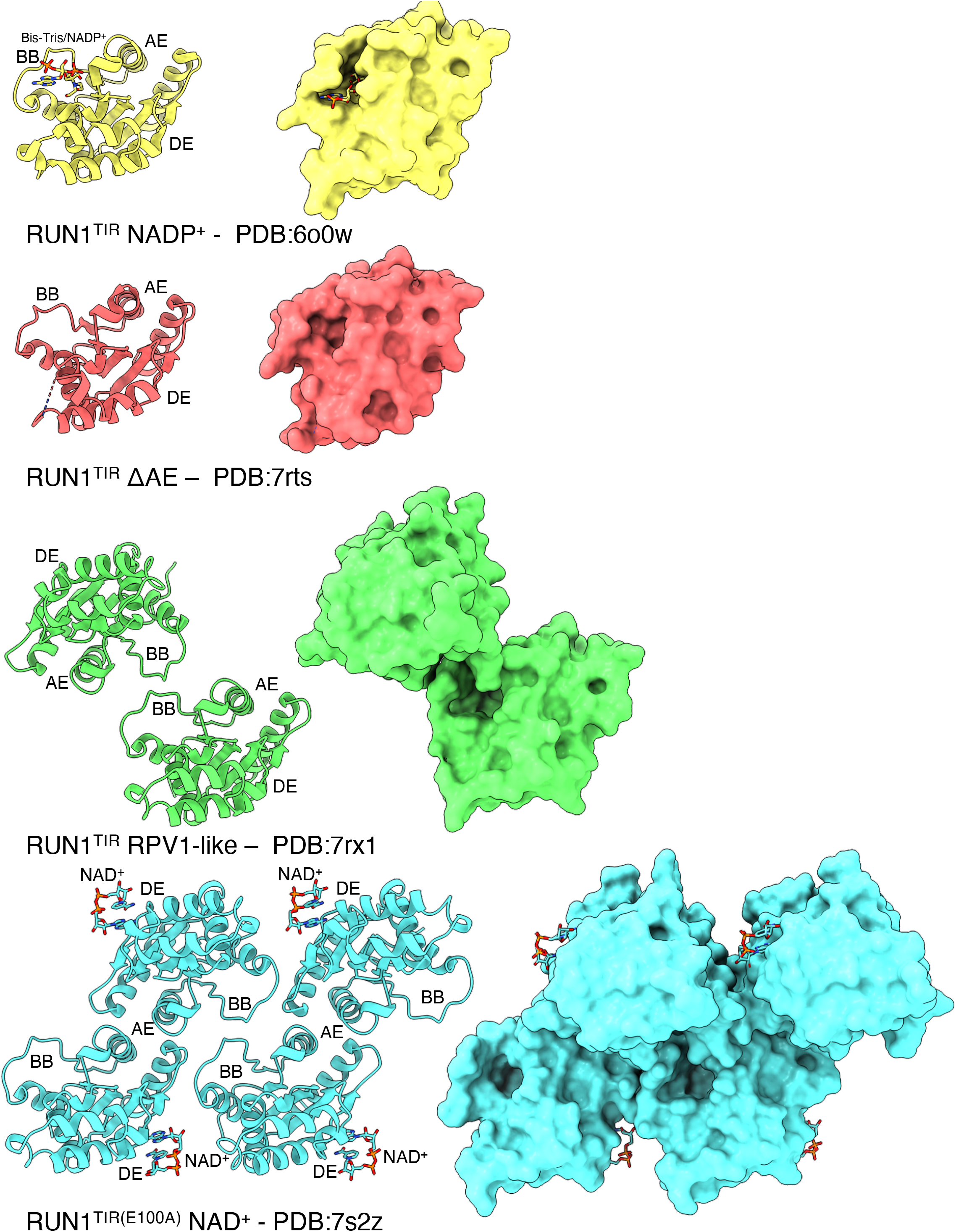
Crystal structures of RUN1^TIR^ domain. RUN1^TIR^ bound to NADP^+^ from Horsefield et al. (2019) (yellow, PDB ID: 6o0w); RUN1^TIR^ with no AE interface (red, PDB ID: 7rts); RUN1^TIR^ with packing arrangement similar to RPV1^TIR^ structure (PDB ID: 5ku7) from Williams et al. (2016) (green, PDB ID: 7rx1); RUN1^TIR(E100A)^ bound to NAD^+^ (cyan, PDB ID: 7s2z).

**Figure S3.**
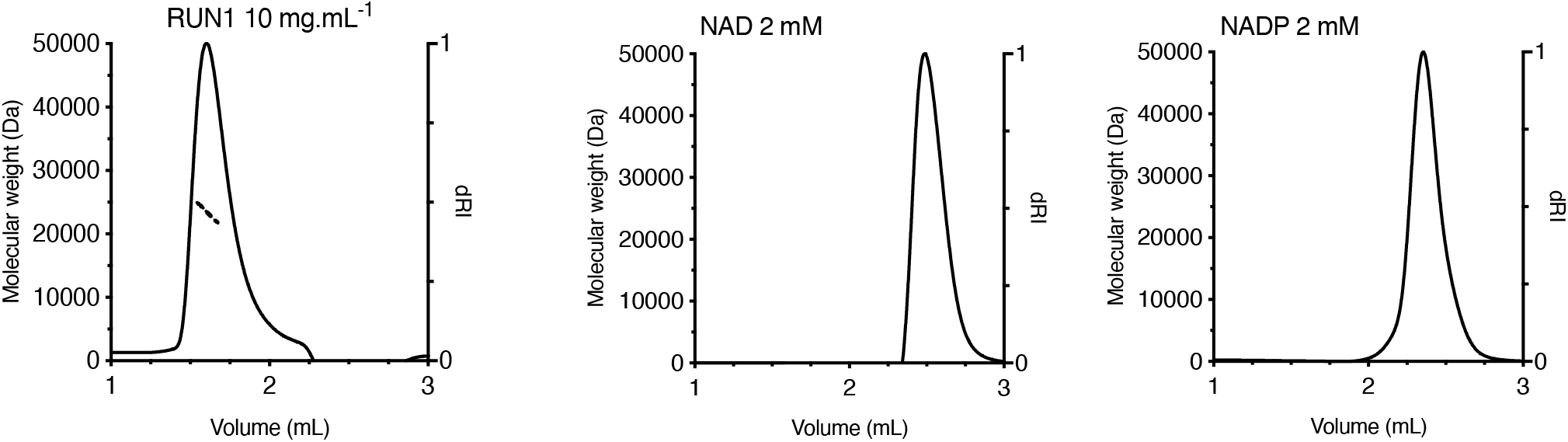
RUN1^TIR^, NAD^+^ and NADP^+^ SEC-MALS traces. The proteins were incubated with 2 mM NAD^+^ or 2 mM NADP^+^ for 20 minutes at room temperature, before SEC-MALS analysis. RUN1^TIR^ 10 mg.mL^-1^ elutes with a peak of a molar mass consistent with a monomeric RUN1^TIR^ (~ 23 kDa). NAD^+^ and NADP^+^ can also be seen on the differential refractive index (dRI) trace, eluting at ~ 2.5 mL. RUN1 in this instance refers to the TIR domain of RUN1, NAD to NAD^+^ and NADP to NADP^+^.

**Figure S4.**
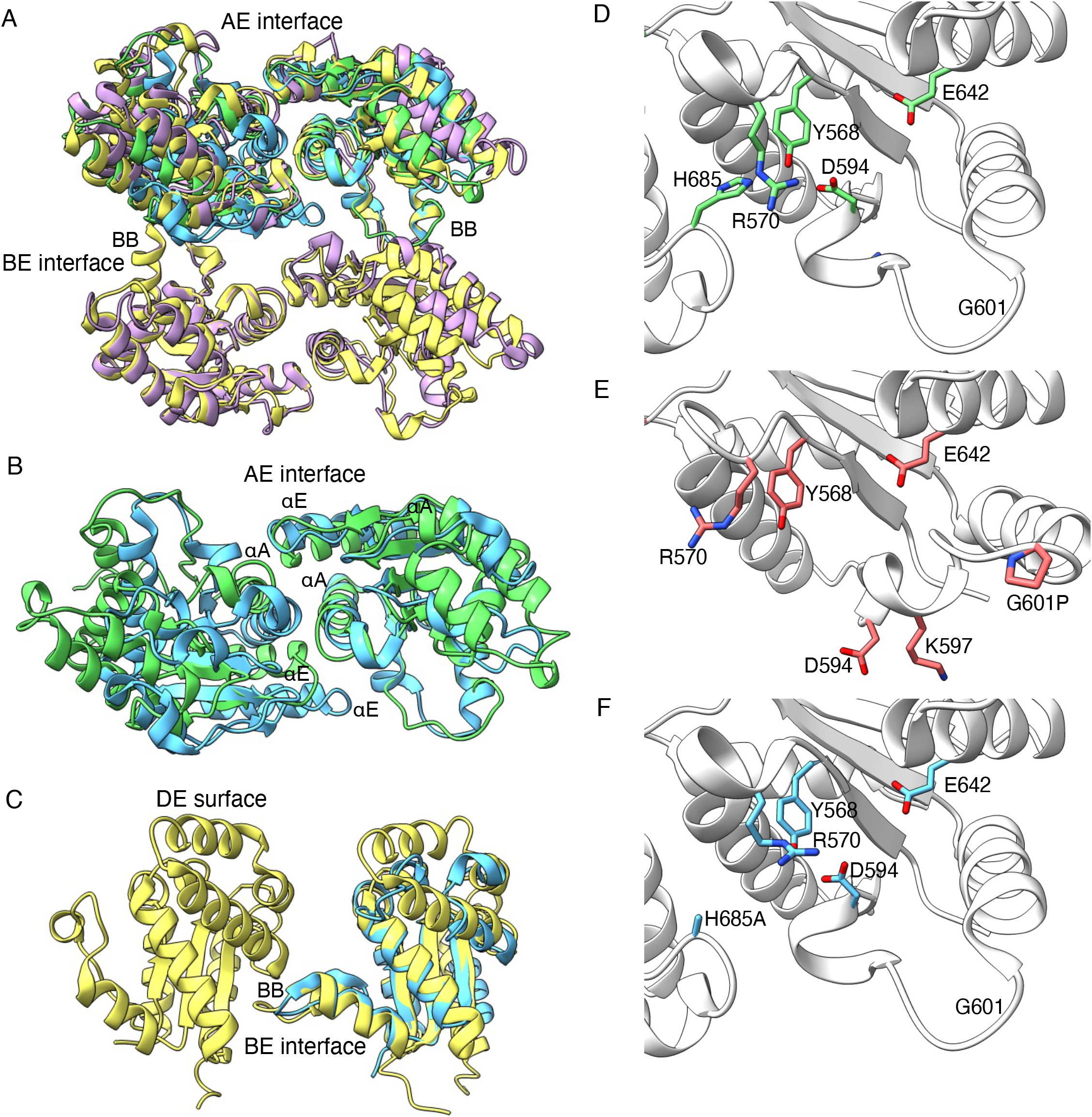
Comparison of the structures of TIR domains from SARM1, RUN1, ROQ1 and RPP1. **A**, superimposition of RPP1^TIR^ (yellow, PDB ID: 7dfv), ROQ1^TIR^ (purple, PDB ID: 7jlx), RUN1^TIR^ (green, PDB ID: 7rx1) and SARM1^TIR^ (cyan, PDB ID: 6o0r), shows a similar packing arrangement between SARM1^TIR^ and plant TIRs. **B**, RUN1^TIR^ AE interface and SARM1^TIR^ interface. **C,** RPP1^TIR^ and SARM1^TIR^ BE interface superimposition. The position of the BB-loop is strikingly similar between RPP1^TIR^ tetramer and SARM1^TIR^. **D-F,** position of residues in the αE helix, and BB-loop and NAD^+^ binding site of SARM1^TIR^ and mutants. **D,** shows the wild type SARM1^TIR^ (PDB ID: 6o0r), **E,** shows SARM1^TIR(G601P)^ (PDB ID: 6o0v). The BB-loop has adopted a closed conformation. **F,** shows SARM1^TIR(H685A)^ (PDB ID: 6o0u), effecting some positions of residues around the NAD^+^ binding site.

## Bibliography

Adams, P. D., Baker, D., Brunger, A. T., Das, R., Dimaio, F., Read, R. J., Richardson, D. C., Richardson, J. S. & Terwilliger, T. C. 2013. Advances, Interactions, and Future Developments in the CNS, Phenix, and Rosetta Structural Biology Software Systems. Annual Review of Biophysics, 42, 265–287.

Beilsten-Edmands, J., Winter, G., Gildea, R., Parkhurst, J., Waterman, D. & Evans, G. 2020. Scaling diffraction data in the DIALS software package: algorithms and new approaches for multi-crystal scaling. Acta Crystallographica Section D, 76, 385–399.

Bernoux, M., Burdett, H., Williams, S. J., Zhang, X., Chen, C., Newell, K., Lawrence, G. J., Kobe, B., Ellis, J. G., Anderson, P. A. & Dodds, P. N. 2016. Comparative analysis of the flax immune receptors L6 and L7 suggests an equilibrium-based switch activation model. Plant Cell, 28, 146–159.

Bernoux, M., Ve, T., Williams, S. J., Warren, C., Hatters, D., Valkov, E., Zhang, X., Ellis, J. G., Kobe, B. & Dodds, P. N. 2011. Structural and functional analysis of a plant resistance protein TIR domain reveals interfaces for self-association, signaling, and autoregulation. Cell Host Microbe, 9, 200–211.

Bi, G., Su, M., Li, N., Liang, Y., Dang, S., Xu, J., Hu, M., Wang, J., Zou, M., Deng, Y., Li, Q., Huang, S., Li, J., Chai, J., He, K., Chen, Y.-H. & Zhou, J.-M. 2021. The ZAR1 resistosome is a calcium-permeable channel triggering plant immune signaling. Cell, 184, 3528–3541.e12.

Burdett, H., Bentham, A. R., Williams, S. J., Dodds, P. N., Anderson, P. A., Banfield, M. J. & Kobe, B. 2019. The Plant “Resistosome”: Structural Insights into Immune Signaling. Cell Host & Microbe, 26, 193–201.

Chen, V. B., Arendall, W. B., 3rd, Headd, J. J., Keedy, D. A., Immormino, R. M., Kapral, G. J., Murray, L. W., Richardson, J. S. & Richardson, D. C. 2010. MolProbity: all-atom structure validation for macromolecular crystallography. Acta Crystallographica. Section D: Biological Crystallography, 66, 12–21.

Dinesh-Kumar, S. P., Tham, W. H. & Baker, B. J. 2000. Structure-function analysis of the tobacco mosaic virus resistance gene *N*. Proceedings of the National Academy of Sciences of the United States of America, 97, 14789–14794.

Duxbury, Z., Wang, S., Mackenzie, C. I., Tenthorey, J. L., Zhang, X., Huh, S. U., Hu, L., Hill, L., Ngou, P. M., Ding, P., Chen, J., Ma, Y., Guo, H., Castel, B., Moschou, P. N., Bernoux, M., Dodds, P. N., Vance, R. E. & Jones, J. D. G. 2020. Induced proximity of a TIR signaling domain on a plant-mammalian NLR chimera activates defense in plants. Proceedings of the National Academy of Sciences, 117, 18832–18839.

Emsley, P., Lohkamp, B., Scott, W. G. & Cowtan, K. 2010. Features and development of Coot. Acta Crystallographica. Section D: Biological Crystallography, 66, 486–501.

Evans, P. 2006. Scaling and assessment of data quality. Acta Crystallographica Section D, 62, 72–82.

Goddard, T. D., Huang, C. C., Meng, E. C., Pettersen, E. F., Couch, G. S., Morris, J. H. & Ferrin, T. E. 2018. UCSF ChimeraX: Meeting modern challenges in visualization and analysis. Protein Science, 27, 14–25.

Horsefield, S., Burdett, H., Zhang, X., Manik, M. K., Shi, Y., Chen, J., Qi, T., Gilley, J., Lai, J.-S., Rank, M. X., Casey, L. W., Gu, W., Ericsson, D. J., Foley, G., Hughes, R. O., Bosanac, T., Von Itzstein, M., Rathjen, J. P., Nanson, J. D., Boden, M., Dry, I. B., Williams, S. J., Staskawicz, B. J., Coleman, M. P., Ve, T., Dodds, P. N. & Kobe, B. 2019. NAD+ cleavage activity by animal and plant TIR domains in cell death pathways. Science, 365, 793–799.

Joseph, A. P., Srinivasan, N. & De Brevern, A. G. 2012. Cis-trans peptide variations in structurally similar proteins. Amino Acids, 43, 1369–1381.

Kabsch, W. 2010. XDS. Acta Crystallographica Section D, 66, 125–132.

Krissinel, E. & Henrick, K. 2007. Inference of Macromolecular Assemblies from Crystalline State. Journal of Molecular Biology, 372, 774–797.

Ma, S., Lapin, D., Liu, L., Sun, Y., Song, W., Zhang, X., Logemann, E., Yu, D., Wang, J., Jirschitzka, J., Han, Z., Schulze-Lefert, P., Parker, J. E. & Chai, J. 2020. Direct pathogen-induced assembly of an NLR immune receptor complex to form a holoenzyme. Science, 370, eabe3069.

Martin, R., Qi, T., Zhang, H., Liu, F., King, M., Toth, C., Nogales, E. & Staskawicz, B. J. 2020. Structure of the activated ROQ1 resistosome directly recognizing the pathogen effector XopQ. Science, 370, eabd9993.

Mccoy, A. J. 2007. Solving structures of protein complexes by molecular replacement with Phaser. Acta Crystallographica. Section D: Biological Crystallography, 63, 32–41.

Mestre, P. & Baulcombe, D. C. 2006. Elicitor-mediated oligomerization of the tobacco N disease resistance protein. Plant Cell, 18, 491–501.

Nishimura, M. T., Anderson, R. G., Cherkis, K. A., Law, T. F., Liu, Q. L., Machius, M., Nimchuk, Z. L., Yang, L., Chung, E.-H., El Kasmi, F., Hyunh, M., Osborne Nishimura, E., Sondek, J. E. & Dangl, J. L. 2017. TIR-only protein RBA1 recognizes a pathogen effector to regulate cell death in Arabidopsis. Proceedings of the National Academy of Sciences of the United States of America, 114, E2053–E2062.

Pergolizzi, G., Butt, J. N., Bowater, R. P. & Wagner, G. K. 2011. A novel fluorescent probe for NAD-consuming enzymes. Chemical Communications, 47, 12655–12657.

Schreiber, K. J., Bentham, A., Williams, S. J., Kobe, B. & Staskawicz, B. J. 2016. Multiple domain associations within the Arabidopsis immune receptor RPP1 regulate the activation of programmed cell death. PLOS Pathogens, 12, e1005769.

Studier, F. W. 2005. Protein production by auto-induction in high density shaking cultures. Protein Expression and Purification, 41, 207–234.

Swiderski, M. R., Birker, D. & Jones, J. D. 2009. The TIR domain of TIR-NB-LRR resistance proteins is a signaling domain involved in cell death induction. Molecular Plant-Microbe Interactions, 22, 157–165.

Tameling, W. I., Elzinga, S. D., Darmin, P. S., Vossen, J. H., Takken, F. L., Haring, M. A. & Cornelissen, B. J. 2002. The tomato R gene products I-2 and MI-1 are functional ATP binding proteins with ATPase activity. Plant Cell, 14, 2929–2939.

Tameling, W. I. L., Vossen, J. H., Albrecht, M., Lengauer, T., Berden, J. A., Haring, M. A., Cornelissen, B. J. C. & Takken, F. L. W. 2006. Mutations in the NB-ARC domain of I-2 that impair ATP hydrolysis cause autoactivation. Plant Physiology, 140, 1233–1245.

Wan, L., Essuman, K., Anderson, R. G., Sasaki, Y., Monteiro, F., Chung, E.-H., Osborne Nishimura, E., Diantonio, A., Milbrandt, J., Dangl, J. L. & Nishimura, M. T. 2019. TIR domains of plant immune receptors are NAD+-cleaving enzymes that promote cell death. Science, 365, 799–803.

Wang, J., Hu, M., Wang, J., Qi, J., Han, Z., Wang, G., Qi, Y., Wang, H.-W., Zhou, J.-M. & Chai, J. 2019a. Reconstitution and structure of a plant NLR resistosome conferring immunity. Science, 364, eaav5870.

Wang, J., Wang, J., Hu, M., Wu, S., Qi, J., Wang, G., Han, Z., Qi, Y., Gao, N., Wang, H.-W., Zhou, J.-M. & Chai, J. 2019b. Ligand-triggered allosteric ADP release primes a plant NLR complex. Science, 364, eaav5868.

Williams, S. J., Sohn, K. H., Wan, L., Bernoux, M., Sarris, P. F., Segonzac, C., Ve, T., Ma, Y., Saucet, S. B., Ericsson, D. J., Casey, L. W., Lonhienne, T., Winzor, D. J., Zhang, X., Coerdt, A., Parker, J. E., Dodds, P. N., Kobe, B. & Jones, J. D. 2014. Structural basis for assembly and function of a heterodimeric plant immune receptor. Science, 344, 299–303.

Williams, S. J., Sornaraj, P., Decourcy-Ireland, E., Menz, R. I., Kobe, B., Ellis, J. G., Dodds, P. N. & Anderson, P. A. 2011. An autoactive mutant of the M flax rust resistance protein has a preference for binding ATP, whereas wild-type M protein binds ADP. Molecular Plant-Microbe Interactions, 24, 897–906.

Williams, S. J., Yin, L., Foley, G., Casey, L. W., Outram, M. A., Ericsson, D. J., Lu, J., Boden, M., Dry, I. B. & Kobe, B. 2016. Structure and function of the TIR domain from the grape NLR protein RPV1. Frontiers in Plant Science, 7, 1850.

Zhang, X., Bernoux, M., Bentham, A. R., Newman, T. E., Ve, T., Casey, L. W., Raaymakers, T. M., Hu, J., Croll, T. I., Schreiber, K. J., Staskawicz, B. J., Anderson, P. A., Sohn, K. H., Williams, S. J., Dodds, P. N. & Kobe, B. 2017. Multiple functional self-association interfaces in plant TIR domains. Proceedings of the National Academy of Sciences of the United States of America, 114, E2046–E2052.

